# Effective Matrix Designs for COVID-19 Group Testing

**DOI:** 10.1101/2022.08.22.504823

**Authors:** David Brust, Johannes J. Brust

## Abstract

**Background:** Grouping samples with low prevalence of positives into pools and testing these pools can achieve considerable savings in testing resources compared with individual testing in the context of COVID-19. We review published pooling matrices, which encode the assignment of samples into pools and describe decoding algorithms, which decode individual samples from pools. Based on the findings we propose new one-round pooling designs with high compression that can efficiently be decoded by combinatorial algorithms. This expands the admissible parameter space for the construction of pooling matrices compared to current methods.

**Results:** By arranging samples in a grid and using polynomials to construct pools, we develop direct formulas for an Algorithm (Polynomial Pools (PP)) to generate assignments of samples into pools. Designs from PP guarantee to correctly decode all samples with up to a specified number of positive samples. PP includes recent combinatorial methods for COVID-19, and enables new constructions that can result in more effective designs.

**Conclusion:** For low prevalences of COVID-19, group tests can save resources when compared to individual testing. Constructions from the recent literature on combinatorial methods have gaps with respect to the designs that are available. We develop a method (PP), which generalizes previous constructions and enables new designs that can be advantageous in various situations.

## Background

The development of vaccines for COVID-19 has constituted a major breakthrough for the return to pre-pandemic life. However, while vaccines are becoming globally widespread, levels of caution prevail. Even with vaccines, monitoring for the evolution of mutations or detecting new outbreaks calls for continued vigilance. Therefore, *testing* is likely to prevail as a vital mechanism to inform decision making in the near future. In order to conserve scarce testing resources, the Centers for Disease Control and Prevention (CDC) in the United States have endorsed so-called group/pooling test methods (likewise, group testing is also being deployed in Central Europe, China, India, Israel and more nations). Such methods date to Dorfman’s early work in 1943 [1], and can be expressed using linear systems. The basic principle underlying pooling tests is the observation that to efficiently detect positive cases among a population with a relatively low occurrence prevalence, it can be advantageous to test groups of samples instead of testing all individual samples.

### One-round Methods

For a set of *N* samples 𝒮 = {*v*_0_, *v*_1_, *v*_2_, …, *v*_*N−*1_}, a basic pooling test pools (mixes) all samples into one group, and subsequently uses one test on this group. If the pool is negative then it can be concluded that each individual sample must be negative. If the pool tests positive at least one sample is positive, and all individuals are retested. This strategy is an example of a 2-round pooling test (in general multi-stage). Based on information from previous testing rounds multi-stage tests update the samples to be tested in the next round. Such methods enable desirable reductions in the number of total needed tests, however they come at the expense of longer times to receive results (since each round adds approximately the same amount of time). An approach that overcomes potentially long wait times is a one-round method [2].

The simultaneous identification of the samples can be represented by a linear system. Since being positive (having the disease) or being negative (not having the disease) are binary states, binary digits i.e., *v* ∈ {0, 1} will be used throughout. Additions and multiplications with *v* are defined by bit-wise OR (+) and AND (∗) operations. Most of the arithmetic carries over expectedly, with the exception that 1 = 1 + 1 in binary form. Suppose now that for certain integers *q* ≥ 2 and *d* ≥ 2 the samples 𝒮 = {*v*_0_, *v*_1_, *v*_2_, …, *v*_*N−*1_} with *N* = *q*^*d*^ have been collected. We would like to infer the states of the elements of vector **v** ∈ {0, 1}^*N*^ based on *M* < *N* observed test results, encoded in vector **w** ∈ {0, 1}^*M*^. Which elements of **v** are included in which test of **w** is determined by the binary matrix **M** ∈ {0, 1}^*M* × *N*^. Specifically, if test *i* contains sample *c* then **M**_*i,c*_ = 1. The Polymerase Chain Reaction (PCR) technology is used to generate the test outcomes in **w** (by amplifying the detectable viral load in each pool). All essential information about the pooling tests is therefore encoded in the linear system

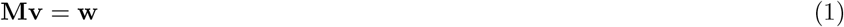

In such schemes pools of size *m* < *N* are formed and simultaneously tested. The distinct composition of pools with respect to their individual samples is essential for one-round methods to exactly identify the true positive samples.

### Pooling Matrix **M**

The distribution of nonzeros in the pooling matrix is equivalent to the design of pooling tests. Importantly, meaningful constructions of **M** enable one to exactly identify up to a certain number of positive samples using only the matrix and the observed values in **w**. Two properties that ensure that up to *k* ≥ 1 positive samples are guaranteed to be identified are:

P.1. Each pair of different samples (*v*_*i*_, *v*_*j*_) occurs at most (*d* − 1) times (i.e., any column pair has at most (*d* − 1) common nonzeros),

P.2. Each sample is contained in *k*(*d* − 1) + 1 tests.

To see the role of these properties we describe the main processes for decoding the observed outcomes to the desired sample values (cf. [3, Corollary 1], too).

### Decoding

Suppose that there are up to *k* positives among all samples. Select and consider an arbitrary sample. If any of the first *k*(*d* − 1) tests with this sample are negative the sample must be negative. Otherwise there are two possible cases. In case one, the sample was misfortunately always paired with one of the *k* positives, even though it is negative. The second case is that the sample is indeed positive. Hence, based on property P.2., if the next test with this sample (i.e., test *k*(*d* − 1) + 1) is negative the sample must be negative. If the test is positive the sample must be positive. By P.2., each sample is contained in *k*(*d* − 1) + 1 tests and therefore all samples can be identified. Even though not absolutely necessary for the decoding of samples, an additional desirable property may be to have uniform pool sizes, since equipment and PCR setups typically have uniform layouts. An effective and fast algorithm to decode the observed **w** into **v** is Combinatorial Orthogonal Matching Pursuit (COMP) [4]. This algorithm declares *v*_*j*_ = 1 if all tests in which *v*_*j*_ is contained are positive and correctly detects up to *k* positives when **M** has properties P.1. and P.2. A straightforward representation of the COMP decoding algorithm is, with “.*” meaning element-wise multiplication:

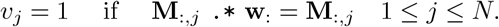

### Illustration

In an example scheme with *d* = 2, *q* = *m* = 4, *N* = *q*^*d*^ = 16 and *M* = 8 the linear system for doing *M* simultaneous tests (where rows are labeled [𝓁_1_-𝓁_8_] and columns [0-15]) is given by:

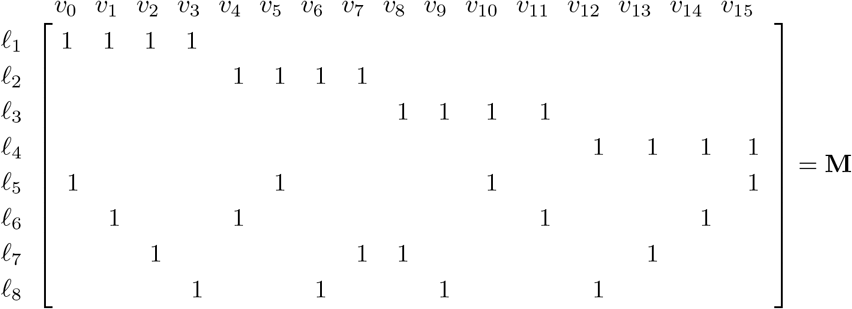

Here, the first row of **M** corresponds to pooling samples (*v*_0_, *v*_1_, *v*_2_, *v*_3_) in test 1 (labeled 𝓁_1_). Observe that *d* − 1 = 1 and that the matrix satisfies property P.1., since each pair of columns has at most one common nonzero. Moreover, note that each sample occurs in *k*(*d* − 1) + 1 = 2 tests since each column has 2 nonzeros (also here *k* = 1). Based on P.1. and P.2., **M** can therefore be used to exactly identify **v** in (1) when **v** contains up to one nonzero element. This design consequently uses 8 tests to identify up to one positive among 16 samples. The “Gain” (*G*) in using **M**, when compared to testing all samples individually, is the ratio: *G* = *N/*(Number of tests) = 16*/*8 = 2.

For the special case that *d* = 2 and when each test has the same number of *m* samples, **M** is called a *multipool* matrix, and *k* + 1 represents the multiplicity of the matrix [5]. Moreover, multipool matrices are a specific *block design* from combinatorial designs [6, Part II]. Concretely, a balanced incomplete block design (BIBD) is a pair (𝒮, ℬ) where 𝒮 is a set of *N* elements and ℬ is a collection of *b m*-subsets of 𝒮 (called blocks) such that each element of 𝒮 is contained in exactly *r* blocks and any 2-subset of 𝒮 is contained in exactly *λ* blocks. Therefore, a mulitpool matrix corresponds to a BIBD with *λ* = 1 and *r* = *k* + 1.

### Recent pooling methods applied to COVID-19

In the following, we review published pooling methods for in infectious disease detection and high-troughput screening. Emphasis is placed on the state of the art literature on pooling methods applied in the context of COVID-19 testing.

The “square array” and related “grid” methods are commonly used for the generation of pooling designs and their evaluation in the presence of testing error [7, 8, 9]. These designs are characterized by arranging the samples in the form of a grid that can be square. Samples are then grouped together by pooling along rows, columns or sloped lines of the grid.

An early study in the context of COVID-19 disease detection proposes a non-adaptive, one round grid design termed “multipool” [5]. The notation 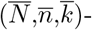multipool is introduced to help distinguish the notation. 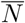 is the number of samples in the pooling scheme, 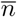 is the order and 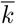 is the “multiplicity”, the number of pools each sample appears in that bounds the maximum number of positives the method guarantees to detect with the COMP algorithm [5]. The multipool design yields pools of size 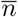, a prime number, capable to test up to 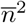 samples in one round of testing. Summarizing, the samples are arranged in an 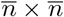 square grid. Pools are formed by pooling along rows, columns or sloped lines of the grid. Thus all samples are covered once by a parallel class [6, I.1.6] of 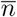 pools corresponding to one pooling direction. The resulting “multipool” consists of the collection of *k* ≤ *m* + 1 parallel classes, capable of detecting up to *k* − 1 positive samples.

For certain situations, an important parameter in the construction of pooling designs is the pool size *m* which determines the number of tests needed in order to detect the expected amount of positives and hence the pooling efficiency [10, 11]. The authors of [10] first describe the dependency of pooling efficiency on the pool size and prevalence of positive samples in the context of two-stage pooling tests. The argument is based on the calculation of the expected value of analyses (tests) per sample as a function of pool size required to identify the anticipated number of positive samples. Derivation of this expected value with respect to the pool size and target prevalence yields a formula for the optimal pool size in their model. Tables with optimum pool sizes for a wide range of expected prevalences are provided for referencing. A pooling design method that is flexible and whose pool size can be tailored to match the optimum at given prevalence is expected to yield higher pooling efficiency.

Pooling designs whose set of pools can be partitioned into subsets which each partition all samples are called resolvable [6, I.1.6]. The so called “Shifted Transversal Design” (STD) [3] is a non-adaptive, combinatorial pool testing method designed for the application in binary high throughput screening and is an example of resolvable designs. A variation of STD, adapted and applied to high-throughput drug screening is presented in [12]. The STD pooling scheme is centered around the parameter *q*, which must be prime. STD yields a set of pools which are grouped into *q* +1 “layers”, each of which in turn partitions all samples thus making it resolvable. The first *q* layers share a regular structure and are therefore called homogeneous. The last remaining layer has a specific construction that differs from the homogeneous layers and is hence called singular. A central characteristic of STD is that pairs of samples should occur together in the same pool only a limited number of times (co-occurrence of samples). In [3] the co-occurrence is denoted by Γ and the relationship for the maximum number of *t* positives that are guaranteed to be discovered by a pooling scheme for a fixed value of *q* is given by Γ*t* + 1 ≤ *k* where *k* is the number of layers. Since STD based on *q* generates up to *q* + 1 layers it holds that Γ*t* ≤ *q*.

A recent study [13] proposes a new algorithm “packing the pencil of lines” (PPoL) for the construction of pooling tests in the context of COVID-19 infection detection. Its conception relies on bipartite graphs and properties of finite projective planes. The construction starts with a bipartite graph whose nodes are connected according to the incidence structure of points and lines prescribed by a finite projective plane of prime power order *m*. The initial graph consists of *m*^2^ + *m* + 1 sample and *m*^2^ + *m* + 1 pool nodes, which correspond to the points and lines of the finite projective plane respectively. Specifically, there is an edge between a sample node and a pool node in the graph, if the corresponding points and lines in the finite projective plane are incident. Trimming the graph by removing nodes and associated edges according to the PPoL algorithm results in a pooling scheme for testing *m*^2^ samples with characteristic parameters (*d*_1_, *d*_2_). It can be represented as a *d*_1_*m*×*m*^2^ binary pooling matrix where the parameters *d*_1_ and *d*_2_ are the number of non-zero elements in each column and row of the pooling matrix, respectively. *d*_1_ corresponds to the number of pools that each sample occurs in and *d*_2_ = *m* is the pool size. The compression gain of this pooling matrix is 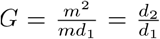 and the number of positives that can be detected is *d*_1_ − 1.

In the same study, the authors also compare the performance of pooling designs generated with PPoL with published approaches from the literature in a probabilistic analysis. Samples are assumed as independent and identically distributed Bernoulli random variables, with a probability of being positive *r*_1_ (prevalence) and negative *r*_0_, respectively. Simulated pooling test results are decoded according to the different pooling matrices using the “two-stage definite defective decoding” (DD2) algorithm [14]. Performance is measured by “expected relative cost” (ERC) which is the expected value of required tests for the detection of positive samples with the DD2 algorithm, divided by the total number of samples in the pooling scheme. In terms of DD2, the ERC is expressed as 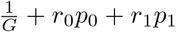, where *p*_0_ and *p*_1_ are the conditional probabilites that a sample cannot be decoded given it is negative and positive, respectively.

PPoL pooling matrices share a common structure and properties with the newly proposed pooling methods from this article, and the performance assessment described above should carry over to the proposed methods in this work.

PPoL [13], P-BEST [15], Tapestry [16, 17] and 2D pooling matrices [18] are compared, where 2D pooling [18] is a particularly simple example of grid designs, which considers pooling along the 8 rows and 12 columns of a standard 96-well plate to form 20 pools in total. The pooling designs in [15, 16, 17] are decoded by compressed sensing algorithms.

It was revealed that pooling schemes that offered the highest performance at low prevalence rates (P-BEST and PPoL-(31,3)) yielded lower performance for increasing prevalence rates. The opposite was also observed where pooling schemes with comparatively lower performance (Tapestry, 2D pooling) did not suffer from higher expected relative cost at higher rates of prevalence. Overall it was revealed that no single design can achieve the highest compression gain for all considered values of prevalence when evaluated with the DD2 algorithm. Instead, the best pooling scheme should be selected depending on the expected prevalence rate of positives.

Summarizing, most of the reviewed pooling designs rely on selecting suitable parameters for the construction. From our observation, limitations in admissible construction parameters for published pooling designs are common. Methods have been proposed, that center around the order parameter *q* and that can test *N* = *q*^*d*^ samples with either *d* = 2 or *d* ≥ 2. Often the order *q* is limited to prime numbers. In cases were *q* was permitted to assume values of prime powers, the constructions were restricted to *d* = 2. This limits the total amount of samples *N* = *q*^2^ that can be tested in one round and thus the maximum achievable compression gain. Published methods often rely on affine planes in the construction of pooling designs restricting the value of the pool size parameter which for affine planes is the same as the order *m* = *q*. As has been demonstrated, the optimal pool size for use in pooling designs depends on the prevalence of positive samples. Designs relying only on fixed values of *m* therefore cannot be adapted to best match the expected prevalence rate.

### Contributions

Addressing the observed limitations in the published methods, we propose novel constructions for one-round combinatorial pooling tests that expand the state of the art methods in three ways. First, the proposed method permits prime power values for the order of the design *q* = *p*^*n*^, which makes the method more flexible with regard to input parameters. Second, it includes pooling matrices based on projective geometries, where the pool size assumes values of *m* = *q* + 1. Because optimal pool size depends on the prevalence of positives [10, 11], projective geometries extend the range of available pool sizes and facilitate matching the design to the expected prevalence ratio. Third, it introduces pooling designs based on prime power order *q* = *p*^*n*^ with dimension *d* ≥ 2 which expands the available parameter space for methods delivering high compression gains for large *N* compared to current methods.

## Results

Our method, PP (Polynomial Pools), is based on arranging samples in a grid and generating pools using polynomials. The degree of the polynomial is *d* − 1 and a sample is represented with a pair of coordinates *v* = (*x, y*). When *d* = 2 our method is analogous to the incidence matrix of an affine plane [6, VII.2.2], [19] and the polynomials reduce to lines. For *p* = *q* a prime number the computations can be done using modulus arithmetic. Specifically, a finite field of order *p* is defined by 𝔽_*p*_ = {0, 1, 2, …}mod *p* and has the important property that arithmetic calculations remain within the field. We label pools by 𝓁 and group samples through the linear relation

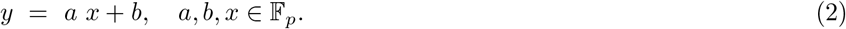

The total number of samples in the affine plane (i.e., *d* = 2) is the square *N* = *p*^2^. For fixed *a, b* in 𝔽_*q*_ and the interval of integers 0 ≤ *i* ≤ *p* − 1 we compute pools using the formulas

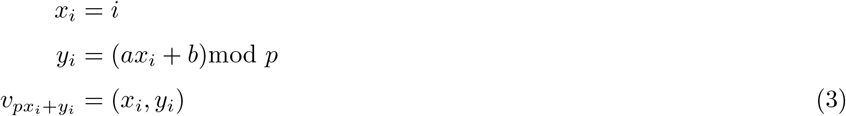

A line/pool containing *p* samples is then the set 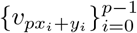. We generalize this method further in three ways. First, for *d* = 2, we include the option of constructing a projective plane [20, VII.2.1]. The projective plane is based on *N* = *p*^2^ + *p* + 1 samples with each pool containing *p* + 1 samples. This makes extended designs possible that use pool sizes, which are not primes. Second, we exploit the fact that affine and projective planes can be generated from prime powers *q* = *p*^*n*^ with *n* ≥ 1. The finite field of order *q* (also known by Galois field) then defines computations. Note that when *p* = *q* (i.e., *n* = 1) and the order of the finite field is prime, then modulus arithmetic similar like in (3) is applicable. However, for prime powers *q* = *p*^*n*^(*n >* 1) the computations in the Galois field use sophisticated techniques. Specifically, the integer elements in the finite field are mapped to polynomials with which arithmetic operations are performed (e.g., addition and multiplication) before being mapped back to integers. Moreover, we include a dimension parameter *d* ≥ 2 which enables designs with high compression gains by generating matrices that grow fast with regard to the total number of samples *N* = (*p*^*n*^)^*d*^.

### Algorithm

To generalize (3), we denote the finite field by 𝔽_*q*_ with order *q* = *p*^*n*^, *n* ≥ 1. Defining indices 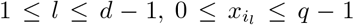 we compute the coordinates of a sample *v* = (*x, y*) using polynomials

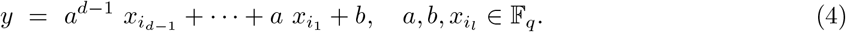

Note that when *d* = 2 and *p* = *q* this reduces to the linear equations from (2). We represent the combination of looping through the 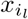 values by a set with *q*^*d−*1^ elements of a sequence of *d* − 1 integers 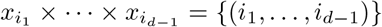, where 0 ≤ *i*_*l*_ ≤ *q* − 1. Without loss of generality, the combination is such that *i*_*d−*1_ cycles every *q* times, *i*_*d−*2_ cycles every *q*^2^ times until *i*_1_ cycles only once. Formulas that compute the sample indices, and thus the corresponding pools, for fixed *a* and *b*, are given by

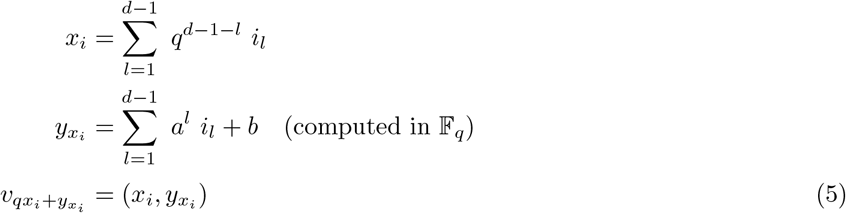

Similar like in (3) computing 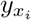 in (5), when *p* = *q* (i.e., prime), can be done using the modulus 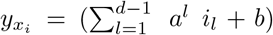mod *q*. Otherwise the additions and multiplications in the expression for 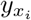 are defined by the operations in the finite field. To illustrate the use of (5) we generate one pool 𝓁 when the slope and intercept are fixed at *a* = 1, *b* = 0 and the parameters are *d* = 3 and *q* = *p* = 3. Table 1 describes the calculations which correspond to the set of samples highlighted in Figure 1.

**Table 1.** Computing a pool, 𝓁 using formula (5) for *d* = 3, *q* = 3, *a* = 1, *b* = 0

| $i_1$ | $i_2$ | $x_i$ | $y_{x_i}$ | $qx_i + y_{x_i}$ | $\ell$ |
| --- | --- | --- | --- | --- | --- |
| 0 | 0 | 0 | 0 | 0 | $v_0$ |
| 0 | 1 | 1 | 1 | 4 | $v_4$ |
| 0 | 2 | 2 | 2 | 8 | $v_8$ |
| 1 | 0 | 3 | 1 | 10 | $v_{10}$ |
| 1 | 1 | 4 | 2 | 14 | $v_{14}$ |
| 1 | 2 | 5 | 0 | 15 | $v_{15}$ |
| 2 | 0 | 6 | 2 | 20 | $v_{20}$ |
| 2 | 1 | 7 | 0 | 21 | $v_{21}$ |
| 2 | 2 | 8 | 1 | 25 | $v_{25}$ |

**Figure 1.**
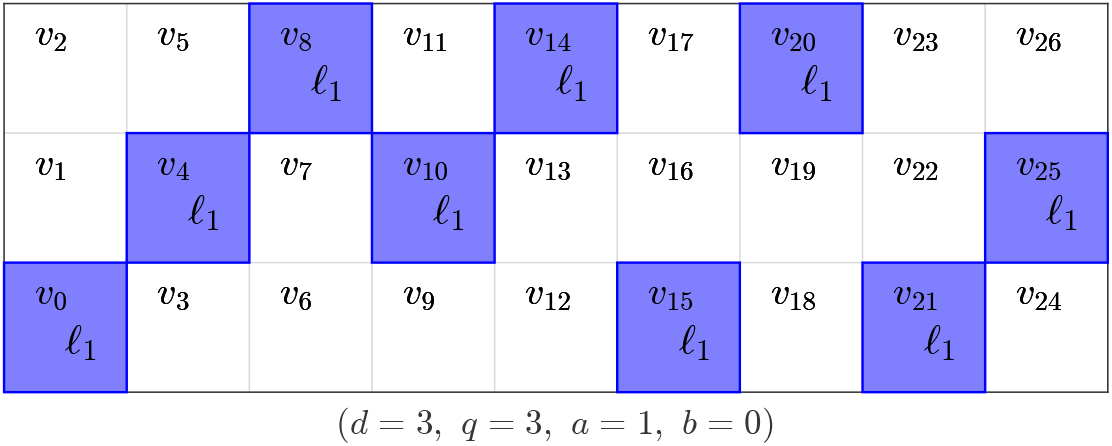
For given parameters, samples are pooled using the polynomial relations from (5). Computation of the indices for samples from the pool is shown in Table 1.

An algorithm that computes sample pools according to polynomial pooling for a desired set of input parameters is given in Algorithm 1. The inputs include *q* = *p*^*n*^, *d* and *k*. The meaning of the parameters is: *order* (*q*) a prime power *q* = *p*^*n*^, *n* ≥ 1, *p* (prime), which includes and extends prime number constructions; *dimension* (*d*) enables scaling to large sample sizes *N* = *q*^*d*^; *characteristic* (*k*) is the limit of positive samples up to which the design exactly decodes all samples. An additional “if else” statement is included, which can be used to generate *q* extra pools when *a* = *q*, i.e., the polynomials have slope of infinity in 𝔽_*q*_. The pools in this situation are formed by grouping *q*^*d−*1^ consecutive samples together. Note that this part of the algorithm is only invoked when the maximum number of positives for the design is requested 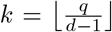. In this case we compute the sample index and define the corresponding point by a different formula.

#### Algorithm 1

(`PP` (`PolynomialPools`))

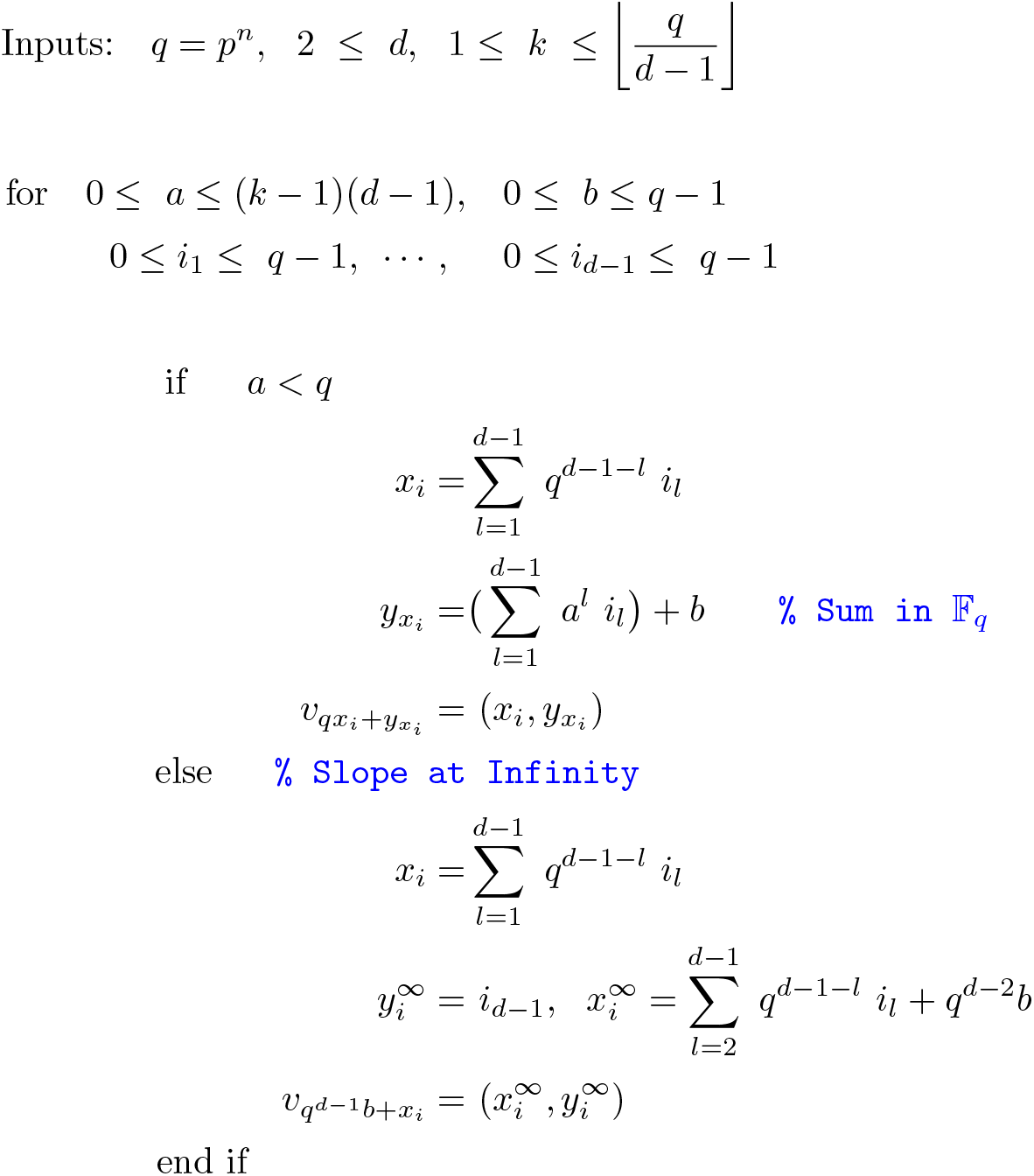

Algorithm 1 generates designs that satisfy properties P.1. and P.2. of a pooling matrix, which guarantee to correctly decode all samples up to *k* positives. A design based on the algorithm is denoted PP(*q, d, k*). Lemmas 1 and 2 relate Algorithm 1 to the relevant properties P.1. and P.2. We prove the Lemmas in the Section “Methods”.

#### Lemma 1

*Any pair of samples v*_*i*_, *v*_*j*_, *for i* ≠ *j from a design of Algorithm 1 occurs at most d* − 1 *times*.

#### Lemma 2

*Each sample v*_*i*_ *is contained in up to k*(*d* − 1) + 1 *tests*.

By satisfying properties P.1. and P.2. the designs from Algorithm 1 guarantee to identify all samples with up to 1 ≤ *k* positives. The size of a pool is the number of elements generated by one polynomial pool 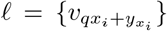, which is *q* = “# elements(𝓁)” (i.e., the number of combinations of *d* − 1 integers 0 ≤ *i*_*l*_ ≤ *q* – 1 with 1 ≤ *l* ≤ *d* − 1). One can also see that designs from PP are resolvable. Figure 2 depicts the *q* − 1 pools that are generated by (5) for a fixed slope *a*, when the intercept ranges through 0 ≤ *b* ≤ *q* − 1. Note that in Figure 2 all *N* samples are pooled into exactly one of the pools, thus forming a resolution. This property comes from the observation that for a fixed slope, changing the intercept results in “parallel” polynomials and leads to “parallel” pools. Furthermore, by varying the parameters *q* and *d* many constructions are feasible, which include the possibility of designs with pool sizes that are for instance not prime.

**Figure 2.**
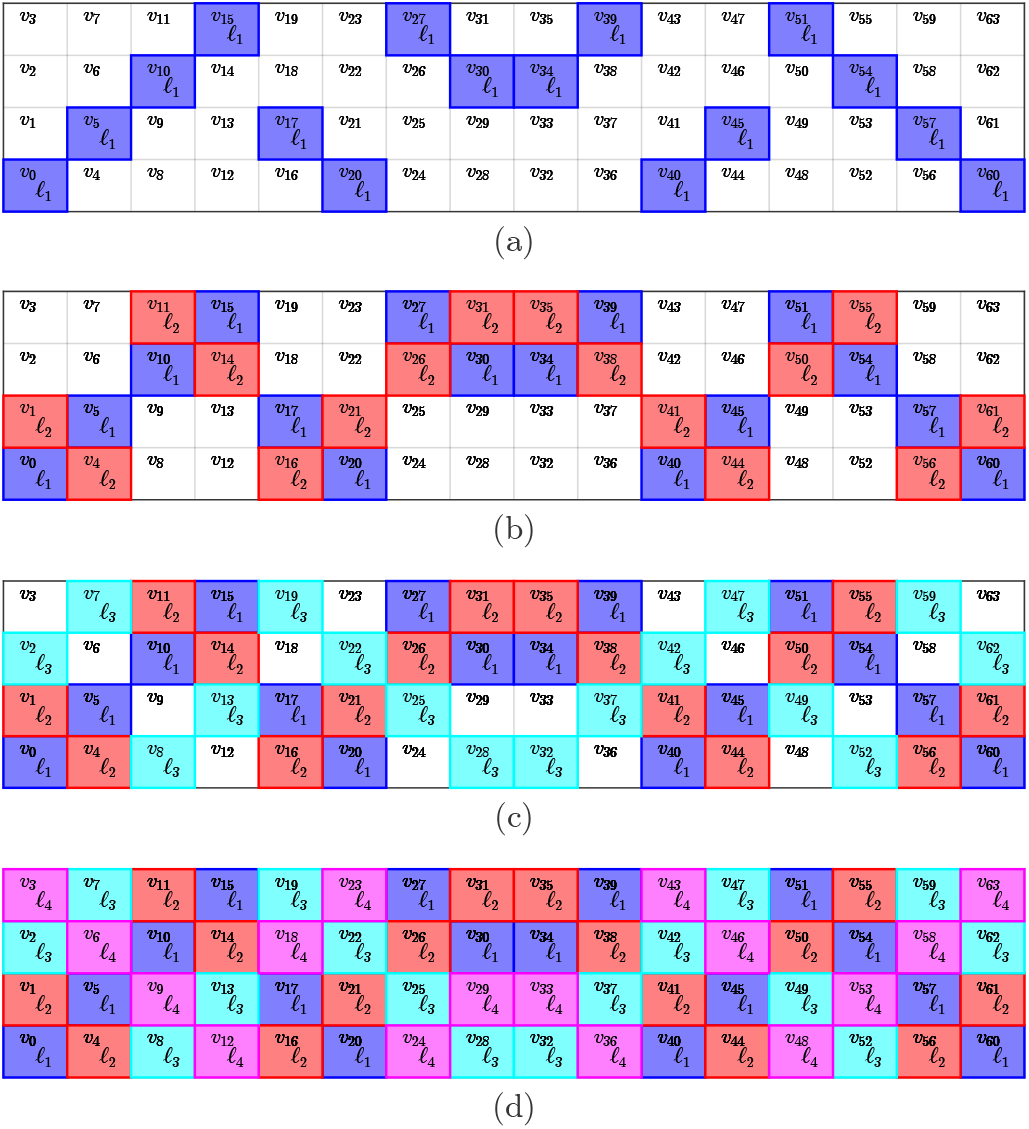
Generating four pools using PP with order *q* = 2^2^ = 4 and slope *a* = 1 when the intercept *b* varies between {0, 1, 2, 3 = *q* − 1} in panels (a),(b),(c),(d).

### Projective Geometry

Furthermore, for a construction from a projective geometry, we modify a design from PP(*q*, 2, *k*). Specifically, we first construct pools from an affine plane of order *q* for 1 ≤ *k* ≤ *q* by invoking PP(*q*, 2, *k*). Then, depending on whether *k* = *q* or *k* < *q* the design is obtained differently. For *k* = *q*, each pool of the affine plane is augmented with one variable. Denote by 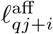 for 1 ≤ *i* ≤ *q* and 0 ≤ *j* ≤ *q* the pools from the affine plane. The projective plane is then obtained by generating the pools 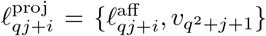 and adding one extra pool 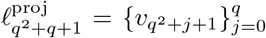. The full projective plane (*k* = *q*) results in a square matrix, which is not advantageous for group testing. Therefore, we next develop an approach for *k* < *q* which is practically more relevant. In particular, the first *q*(*k* + 1) pools are extended with one sample like before, 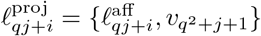 for 1 ≤ *i* ≤ *q* and 0 ≤ *j* ≤ *k*. Then we add *q* – *k* “singleton” pools each containing only one sample so that 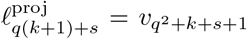 with 1 ≤ *s* ≤ *q* − *k*. A projective geometry enables designs with pool sizes *m* = *q* + 1 (and individual tests), and thus opens up new constructions. We denote designs from projective geometries by PP_PG_(*q*, 2, *k*).

In the following we highlight characteristics of the PP method proposed in this work by comparing it to previously published methods presented in Section “Recent pooling methods applied to COVID-19”.

### Prime power order

We first compare PP to 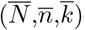-multipool designs [5], where overline notation was introduced in the leading paragraph of Section “Recent pooling methods applied to COVID-19”. The construction of multipools is restricted to prime orders 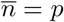. The remaining design parameters follow as 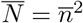 and 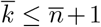. In the particular case of non-prime orders 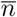, the method is restricted to 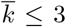. This limits the number of detectable positives to 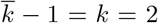 when non-prime orders are chosen. Table 2 compares the pooling designs that can be generated with the 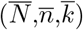-multipool method and with PP of non-prime order *n* = 8 and *N* = 64 for the detection of *k* positives (this set of parameters was proposed in [5, Section 4])

**Table 2.** Pooling desings of non-prime order

| k | multipools [5]<br>desc. | M | PP (this work)<br>desc. | M |
| --- | --- | --- | --- | --- |
| 1 | (64,8,2)-multipool | 16 | PP(8,2,1) | 16 |
| 2 | (64,8,3)-multipool | 24 | PP(8,2,2) | 24 |
| 3 | n.a. | n.a. | PP(8,2,3) | 32 |
| 4 | n.a. | n.a. | PP(8,2,4) | 40 |
| 5 | n.a. | n.a. | PP(8,2,5) | 48 |
| 6 | n.a. | n.a. | PP(8,2,6) | 56 |
| 7 | n.a. | n.a. | PP(8,2,7) | 64 |
| 8 | n.a. | n.a. | PP(8,2,8) | 72 |

Comparison of pooling designs with order $q = \bar{n} = 8$ from multipool [5] and PP methods to test $N = \bar{N} = 64$ samples. The designs from the multipool method can detect up to $k = 2$ positives, whereas PP can detect up to $k = 8$ positives.

For the case considered in Table 2 the multipool method can generate designs for the detection of at most 2 positives whereas PP allows the construction of designs with *k* ≤ *q*. Here we took advantage of the fact that PP can generate designs with prime power orders *q* = *p*^*n*^, *n* ≥ 1, in this case *q* = 2^3^ = 8. We note that designs PP(8,2,*k*), *k* ∈ {7, 8} are listed for completeness and do not have practical significance since the compression gain of these designs is not higher than individual testing.

### Pool sizes and projective planes

Published methods often rely on the affine plane with pool size equivalent to the order of the affine plane *m* = *q* [5, 13]. In the context of pool testing with two-stage decoding algorithms it has been suggested that pool size for optimal pooling efficiency depends on the prevalence [10, 11]. Table 3 shows pool size and corresponding prevalence values in the first two columns as obtained in [10]. Prevalences are defined by maximum detectable positives *k* divided by total samples *N*. Due to the discrete nature of pooling designs and their parameters, it is not always possible to match prescribed values of prevalence exactly, therefore the values of prevalence shown in Table 3 are within ±5% of the reported values. Designs based on projective geometries PP_PG_(*q*,2,*k*) exhibit pool size *m* = *q* + 1. As is apparent from Table 3 the projective geometries complement the PP designs with respect to the covered pool sizes. To the best of our knowledge, the combination of PP and PP_PG_ methods can yield a set of new designs not covered by previously published methods.

**Table 3.** Designs with optimal pool size for various prevalences

| m | prev. | N | M | G | desc. |
| --- | --- | --- | --- | --- | --- |
| 3 | 0.14 | 7 | 5 | 1.4 | PP<sub>PG</sub>(2,2,1) (1 singl.)<sup>†</sup> |
| 4 | 0.071 | 14 | 8 | 1.8 | PP(4,2,1) |
| 5 | 0.043 | 23 | 10 | 2.3 | PP(5,2,1) |
| 6 | 0.031 | 32 | 15 | 2.1 | PP<sub>PG</sub>(5,2,1) (5 singl.)<sup>†</sup> |
| 7 | 0.024 | 42 | 14 | 3 | PP(7,2,1) |
| 8 | 0.017 | 58 | 16 | 3.6 | PP(8,2,1) |
| 9 | 0.015 | 66 | 16 | 4.1 | PP<sub>PG</sub>(8,2,1) |
| 10 | 0.01 | 96 | 31 | 3.1 | PP<sub>PG</sub>(9,2,1) (13 singl.)<sup>†</sup> |
| 11 | 0.0083 | 121 | 22 | 5.5 | PP(11,2,1) |
| 12 | 0.0073 | 137 | 36 | 3.8 | PP<sub>PG</sub>(11,2,1) (14 singl.)<sup>†</sup> |
| 13 | 0.0059 | 169 | 26 | 6.5 | PP(13,2,1) |
| 16 | 0.0039 | 256 | 32 | 8 | PP(16,2,1) |
| 18 | 0.0031 | 318 | 61 | 5.2 | PP<sub>PG</sub>(17,2,1) (27 singl.)<sup>†</sup> |
| 32 | 0.00098 | 1024 | 64 | 16 | PP(32,2,1) |

<sup>†</sup>Number of singleton pools added to projective geometry. This ensures that each sample is tested in the case $k < q$ as described in section “Projective Geometry”. Optimal pool sizes and prevalences are from [10].

### Prime power orders and higher dimensions

We refer to the paragraph of Algorithm 1 in Section “Algorithm”, where we describe the dimension parameter *d*. There we highlight that designs with *d* ≥ 2 yield high compression gains by generating matrices that grow fast with regard to the total number of samples *N* = *q*^*d*^. Of the recently published methods, only STD [3] as well as PP, proposed in this work, take advantage of the exponential growth in total sample number with dimension. Thus STD and PP are able to achieve significantly larger compression gains, especially at low rates of prevalence, than methods that have a fixed dimension of *d* = 2 e.g. PPoL [13] and “multipools” [5]. The significance of the dimension *d* ≥ 2 is illustrated in Table 4 showing design parameters and achieved compression gains of different pooling designs for the detection of 2 positive among 961 samples. These values were proposed for the PPoL-(31,3) design in [13] for prevalences below 2%.

**Table 4.** Pooling designs from published methods

| q | d | M | G | desc. |
| --- | --- | --- | --- | --- |
| 7 | 4 | 49 | 20 | PP(7,4,2) |
| 7 | 4 | 49 | 20 | STD(961,4,7) [3] |
| 31 | 2 | 93 | 10 | PPoL-(31,3) [13] |
| 31 | 2 | 93 | 10 | (961,31,3)-multipools [5] |

Parameters of pooling designs from published methods and from PP (this work) for guaranteed detection of $k = 2$ positives among $N = 961$ samples.

Table 4 lists the designs from PP (this work), STD [3], PPoL [13] and multipools [5] with the highest compression gain for these parameters. In this setting, designs generated by PP and STD are equivalent, the same applies to the designs generated by PPoL and multipools. The main difference to note is the fact that PPoL and multipools are restricted to dimension *d* = 2 and consequently achieve lower compression gains that are, for this set of paramters, half of the compressions achieved with STD and PP respectively.

Higher dimensions *d* ≥ 2 allow the construction of designs with very large *N*. As has been discussed before, the only recently published method that takes advantage of variable dimension is STD [3]. However, STD is restricted to prime orders *q* = *p*. The PP method presented in this work permits construction based on prime order as well as prime power order. Therefore all designs that can be constructed by STD also can be constructed by PP. In the following, examples of designs for large values of *N* are given where application of variable dimension yields large compression gains, especially for low prevalences. Table 5 shows designs for numbers of samples 1000 ≤ *N* ≤ 25000 for the detection of 1 ≤ *k* ≤ 50 positives. Prime power orders *q* = *p*^*n*^, *n* ≥ 1 are highlighted with an asterisk, marking designs that are now available through the PP method proposed in this work.

**Table 5.**
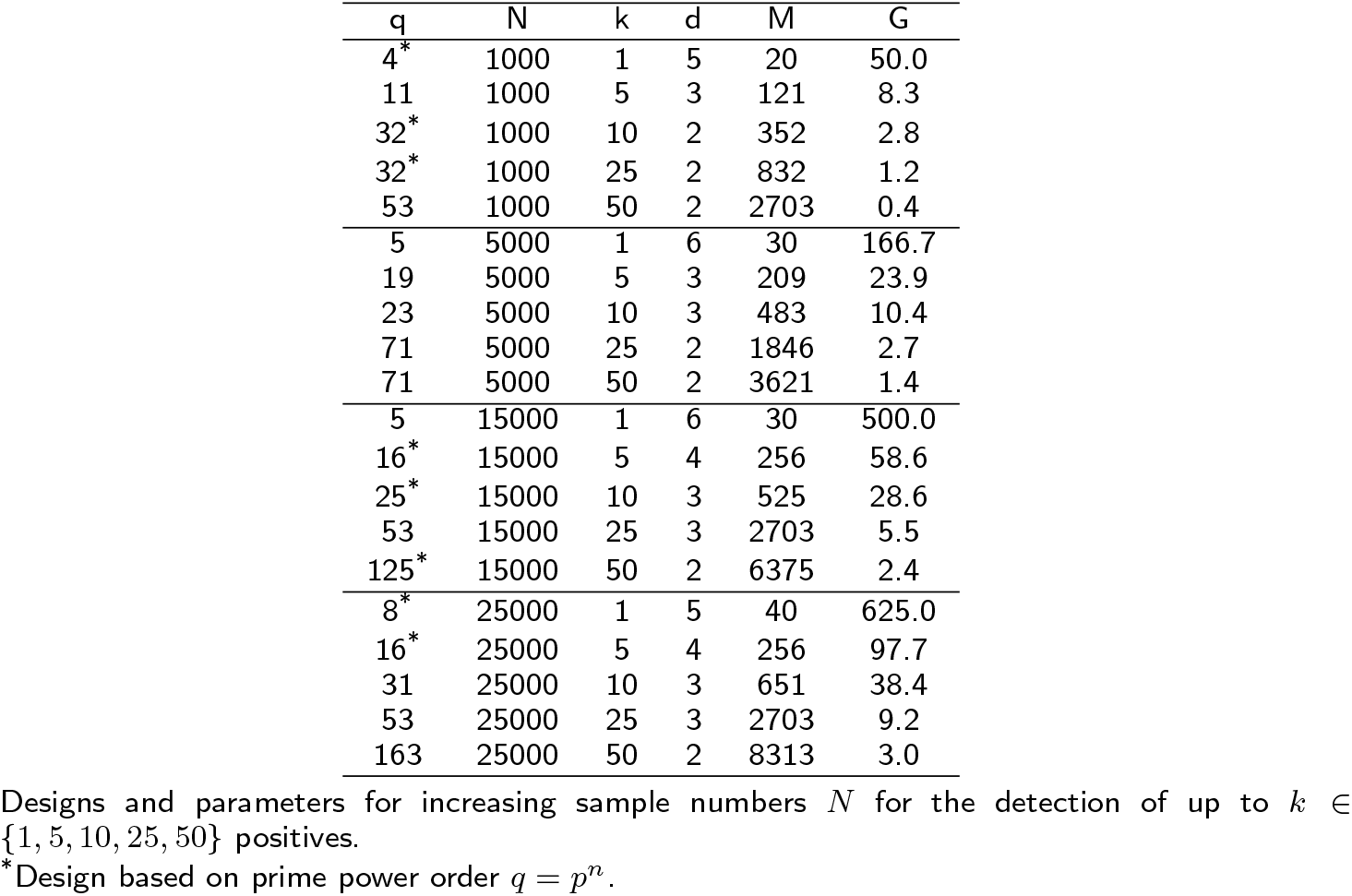
Parameters of large PP designs

| q | N | k | d | M | G |
| --- | --- | --- | --- | --- | --- |
| 4* | 1000 | 1 | 5 | 20 | 50.0 |
| 11 | 1000 | 5 | 3 | 121 | 8.3 |
| 32* | 1000 | 10 | 2 | 352 | 2.8 |
| 32* | 1000 | 25 | 2 | 832 | 1.2 |
| 53 | 1000 | 50 | 2 | 2703 | 0.4 |
| 5 | 5000 | 1 | 6 | 30 | 166.7 |
| 19 | 5000 | 5 | 3 | 209 | 23.9 |
| 23 | 5000 | 10 | 3 | 483 | 10.4 |
| 71 | 5000 | 25 | 2 | 1846 | 2.7 |
| 71 | 5000 | 50 | 2 | 3621 | 1.4 |
| 5 | 15000 | 1 | 6 | 30 | 500.0 |
| 16* | 15000 | 5 | 4 | 256 | 58.6 |
| 25* | 15000 | 10 | 3 | 525 | 28.6 |
| 53 | 15000 | 25 | 3 | 2703 | 5.5 |
| 125* | 15000 | 50 | 2 | 6375 | 2.4 |
| 8* | 25000 | 1 | 5 | 40 | 625.0 |
| 16* | 25000 | 5 | 4 | 256 | 97.7 |
| 31 | 25000 | 10 | 3 | 651 | 38.4 |
| 53 | 25000 | 25 | 3 | 2703 | 9.2 |
| 163 | 25000 | 50 | 2 | 8313 | 3.0 |

Designs and parameters for increasing sample numbers $N$ for the detection of up to $k \in \{1, 5, 10, 25, 50\}$ positives.
\*Design based on prime power order $q = p^n$ .

For each set of parameters in Table 5 the design with the largest compression gain is shown. In general, large compression gains can be observed, especially when few positives are to be detected. Designs with prime power order *q* = *p*^*n*^ are highlighted. We want to emphasize that these achieve the highest compression gain for this parameter set, surpassing designs based on prime order *q* = *p*.

## Discussion

From the previous section (“Results”) it is evident that PP enables designs that have not been possible with existing methods. For instance, multipool matrices for up to 2 positive samples with pool size *m* = 8 and *N* = 64 were suggested in [5]. However, when e.g., up to 3 positives are needed, multipool matrices are not applicable. Nevertheless, with the same pool and sample size PP enables designs for situations with up to 8 positive samples (cf. Table 2). This is reminiscent of the fact that PP permits prime number or prime power orders *q*.

Even though prime power orders are in principle also possible in [13], the methods therein are restricted to *d* = 2 and to affine geometries. PP enables constructions with *d* ≥ 2, which can be useful for large scale testing because of very high compression gains.

For smaller, specifically targeted pool sizes, the option of using projective geometries, PP_PG_, makes it possible to find one-round designs for optimal pool sizes from a two-round method [10] (cf. Table 3)

Finally, PP enables the combination of prime power orders with 2 ≤ *d* and extends high throughput methods with 2 ≤ *d*, but which are constrained to prime number orders [3] (cf. Table 5).

We describe some additional practical considerations. When a certain fixed number of samples 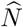 are to be tested, designs may not exist that exactly match 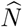. It is then common to construct pooling tests that exceed this number, i.e., 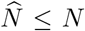, and to “use/tag” the excess samples 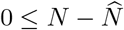 to be known negatives.

Another special situation arises with projective geometries. Since PP_PG_ is obtained by augmenting a subset of *qk* + 1 pools from an affine plane with one sample and by adding a selection of “singleton” tests, the number of samples in tests is twofold: There are *q*(*k* + 1) tests with *q* + 1 samples and *q* − *k* tests with one sample.

Moreover, the PP designs are analyzed in the setting of one-round tests. Embedding pooling matrices with *d* = 2 in a two round setting results in different properties and trade-offs [13]. Future work can be based on the larger class of matrix designs from PP in two-stage pooling test settings.

For finite field computations with prime power order we use the libraries [21, 22].

## Conclusions

Since prevalences or testing settings may change with respect to COVID-19 it is important to have improved options for appropriate test designs. We propose a combinatorial method, Polynomial Pools (PP), that includes constructions from existing approaches and enables new designs. Because of this, PP is at least as effective as existing approaches, but additionally enables designs with more advantages.

## Methods

To motivate the proof for Algorithm 1 (i.e., Lemmas 1,2) we use two basic principles:

- Each 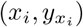 represents one sample
- Each sample and therefore pairs of samples 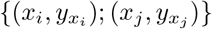 are computed using different polynomial slopes and intercepts (adapted for the “infinity” slope)

Suppose that 1 ≤ *k* < *p* and let *j* ≠ *i*. Select an arbitrary pair of samples 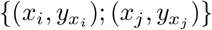, computed using, say, *a* and *b*. Consider the different slope *ā* ≠ *a* and intercept *<u>b</u>* ≠ *b*. There are four possibilities to generate pools in PP, and consequently pairs of samples other than a base pair

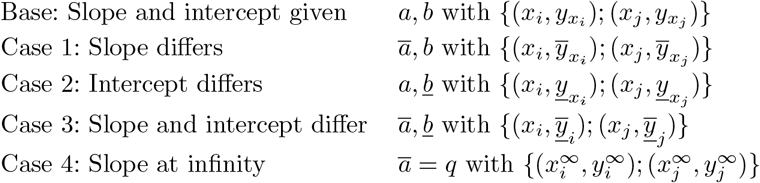

Let 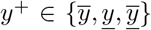 represent a *y* value from one of the Cases 1–3. Note that the conditions for having up to *d* − 1 occurrences of a pair are for *k* ∈ {*i, j*}, *t* ∈ {*i, j*}, *k* ≠ *t*

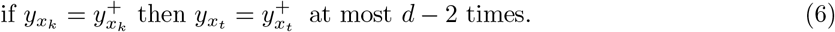

Recall the statement for Lemma 1: *Any pair of samples v*_*i*_, *v*_*j*_, *for i* ≠ *j from a design of Algorithm 1 occurs at most d* − 1 *times*.

*Proof* Using the four cases with 2 ≤ *d* and *q* = *p*^*n*^, 1 ≤ *n*, points (and thus sample indices) are computed by polynomials over the finite field 𝔽_*q*_. In the first three cases the slopes are not “infinite” in F_*q*_ and points are computed using

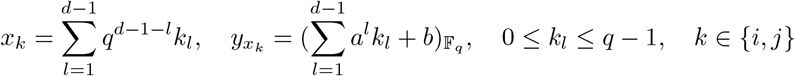

In the fourth case we describe the pools with slope of “infinity”. Suppose that *i* ≠ *j*. Case 1: In this case the slopes differ only. Therefore, *a* ≠ *ā* and there is an integer *g* such that *a*+*g* = *ā* Similarly, there are integers *k* ≠ *t* ∈ {*i, j*} and “subindices” *k*_*l*_, *t*_*l*_ for 1 ≤ *l* ≤ *d* − 1 so that *k*_*l*_ + *g*_*l*_ = *t*_*l*_. Condition (6) is in this case for *k* ≠ *t* ∈ {*i, j*}

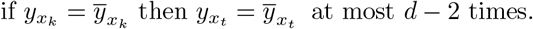

The difference 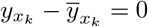 for *k* ∈ {*i, j*} and *k*_*l*_ + *g*_*l*_ = *t*_*l*_ yields

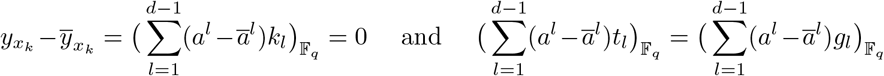

Note that from *ā* = *a* + *g* it holds that *a*^*l*^ − *ā* ^*l*^ = *a*^*l*^ − (*a* + *g*)^*l*^. Thus, *a*^*l*^ − *ā* ^*l*^ is a polynomial of degree at most *l* − 1 with respect to *a* since the first term in the binomial expansion of (*a* − *g*)^*l*^ is *a*^*l*^. Then

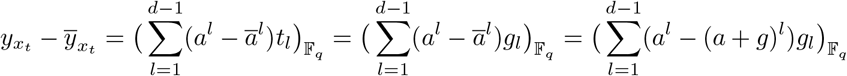

The last equality is a polynomial with degree up to *d* − 2 in F_*q*_ in terms of *a*. Therefore, for *k* ≠ *t* when 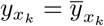 then 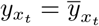 up to *d* − 2 times (at the roots of a polynomial). We subsequently conclude that pairs of samples, including the “base pair” 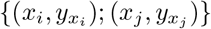 occur up to *d* − 1 times.

Case 2: The intercept differs and with *k* /= *t* ∈ {*i, j*}

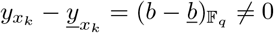

Therefore 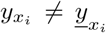 and 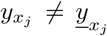 and all samples and corresponding pairs are different in this case.

Case 3: In this case both; slope and intercept differ. Therefore, *a* ≠ *ā b* ≠ *<u>b</u>* and there is an integer *g* such that *a*+*g* = *ā*. As in Case 1 there are integers *k* ≠ *t* ∈ {*i, j*} and “subindices” *k*_*l*_, *t*_*l*_ for 1 ≤ *l* ≤ *d* − 1 so that *k*_*l*_ + *g*_*l*_ = *t*_*l*_. The relevant condition for *k* /= *t* ∈ {*i, j*} is

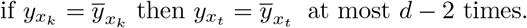

The difference 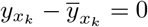 for *k* ∈ {*i, j*} and *k*_*l*_ + *g*_*l*_ = *t*_*l*_ yields

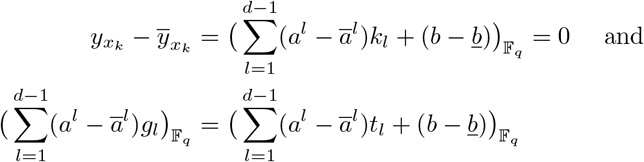

Similar like before note that from *ā* = *a* + *g* it holds that *a*^*l*^ − *ā* ^*l*^ = *a*^*l*^ − (*a* + *g*)^*l*^. Thus, *a*^*l*^ − *ā* ^*l*^ is a polynomial of degree at most *l* − 1 since the first term in the binomial expansion of (*a* − *g*)^*l*^ is *a*^*l*^. Then

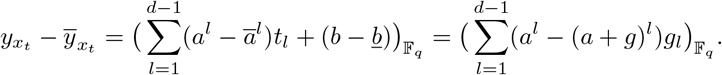

The right hand side in the last equality is a polynomial of degree up to *d* − 2 with respect to *a* in 𝔽_*q*_. Therefore, for *k* ≠ *t* when 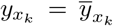 then 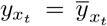 up to *d* − 2 times (at the roots of a polynomial). We subsequently conclude that pairs of samples, including the “base pair” 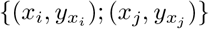 occur up to *d* − 1 times.

Case 4: With the slope at “infinity” in 𝔽_*q*_ the formula for computing points is adapted. Recall that for 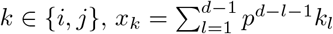 so that the formula for 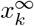 becomes

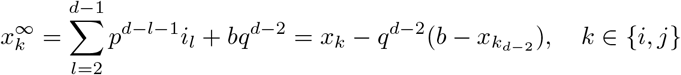

Therefore one sees that for *k* ∈ {*i, j*}

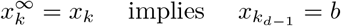

To investigate the condition in (6) we note that

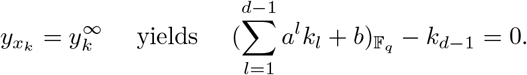

Therefore, for *k* ≠*t* ∈ {*i, j*} (and with *k*_*d−*1_ = *t*_*d−*1_),

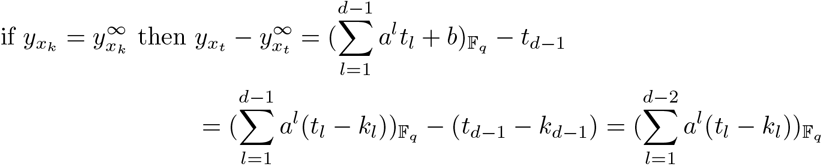

The final equality is a polynomial of degree up to *d* − 2 with respect to *a* and it has up to *d* − 2 roots. Therefore we conclude that a pair of samples occurs up to *d* − 1 times in Case 4. In summary, in all cases with 2 ≤ *d* and *q* = *p*^*n*^, 1 ≤ *n* pairs of samples occur at most *d* − 1 times according with the result of Lemma 1. □

Lemma 2 follows from the previous analysis. Recall the statement of Lemma 2: *Each sample v*_*i*_ *is contained in up to k*(*d* − 1) + 1 *tests*.

*Proof* First note that one pool corresponds to one combination of slope and intercept *a* and *b* (including the infinity slope). When only the intercept *b* differs (and *a* is fixed), each sample occurs exactly once for a combination of *a* and all possible values of *b*. Since 0 ≤ *a* ≤ *k*(*d* − 1) there are *k*(*d* − 1) + 1 different *a* values and thus each sample occurs in *k*(*d* − 1) + 1 tests. □

For *d* = 2 and *q* being a prime number the formulas in PP can be represented using modulus arithmetic and lines. In this situation a further proof of Lemma 1 is given in the “Additional file 1”.

## Supporting information

Supplemental Proof 1

## Appendix

Table 6 contrasts a broader range of published pooling designs with designs using PP. In [13] the PPoL method was compared to the methods from Table 6 excluding PP for varying rates of prevalence up to 5 % in terms of the two-stage decoding algorithm DD2. Performance was measured by expected relative cost (ERC). It was concluded, that there was no single design with the lowest expected relative cost for all rates of prevalence and hence the design should be chosen to match expected prevalence rate. However, according to Figure 11 from [13] the PPoL-(31,3) design has advantages over other published methods as it showed lower ERC at low values of prevalence. At the same time its ERC stayed below competing designs for larger prevalences. Because all PPoL designs are included in PP(*q*,2,*k*) designs with *q* = *p* and further ones are possible with PP we extrapolate that the arguments made in [13] also apply to PP designs. Table 6 illustrates possible advantages with PPoL designs.

**Table 6.**
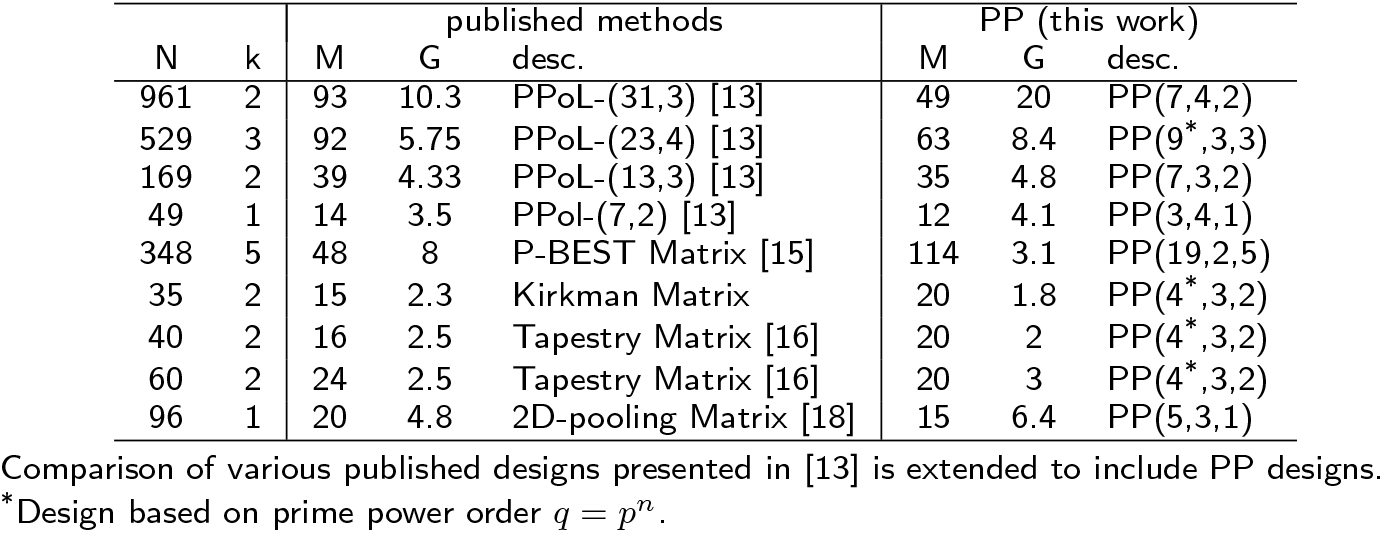
Overview of published methods and corresponing PP designs

| N | k | published methods |  |  | PP (this work) |  |  |
| --- | --- | --- | --- | --- | --- | --- | --- |
|  |  | M | G | desc. | M | G | desc. |
| 961 | 2 | 93 | 10.3 | PPoL-(31,3) [13] | 49 | 20 | PP(7,4,2) |
| 529 | 3 | 92 | 5.75 | PPoL-(23,4) [13] | 63 | 8.4 | PP(9*,3,3) |
| 169 | 2 | 39 | 4.33 | PPoL-(13,3) [13] | 35 | 4.8 | PP(7,3,2) |
| 49 | 1 | 14 | 3.5 | PPoL-(7,2) [13] | 12 | 4.1 | PP(3,4,1) |
| 348 | 5 | 48 | 8 | P-BEST Matrix [15] | 114 | 3.1 | PP(19,2,5) |
| 35 | 2 | 15 | 2.3 | Kirkman Matrix | 20 | 1.8 | PP(4*,3,2) |
| 40 | 2 | 16 | 2.5 | Tapestry Matrix [16] | 20 | 2 | PP(4*,3,2) |
| 60 | 2 | 24 | 2.5 | Tapestry Matrix [16] | 20 | 3 | PP(4*,3,2) |
| 96 | 1 | 20 | 4.8 | 2D-pooling Matrix [18] | 15 | 6.4 | PP(5,3,1) |

Comparison of various published designs presented in [13] is extended to include PP designs.
\*Design based on prime power order $q = p^n$ .

## Acknowledgements

Not applicable.

## Funding

Not applicable.

### Abbreviations

PP: Polynomial Pools
PG: Projective geometry
COMP: Combinatorial Orthogonal Matching Pursuit [4]
STD: Shifted Transversal Design [3]
PPoL: Packing the Pencil of Lines [13]
P-BEST: Pooling-Based Efficient SARS-CoV-2 Testing [15]
DD2: Two-stage Definitive Defectives [14]
ERC: Expected relative cost
PCR: Polymerase Chain Reaction

## Availability of data and materials

The datasets generated and/or analysed during the current study are available in the PP (PolynomialPools) repository, https://github.com/johannesbrust/PP

## Ethics approval and consent to participate

Not applicable.

## Consent for publication

Not applicable.

## Authors’ contributions

DB produced tables 2–6 and implemented a Julia version of the proposed method. JJB derived the polynomial representation of pools and implemented a Matlab version of the proposed method. Both authors were major contributors in writing the manuscript.

## Authors’ information

DB is an engineer with more than 8 years of programming experience. JJB is an applied mathematician who developed affine and projective geometries for COVID-19 pool testing.

## Additional Files

Additional file 1 — Proof of Lemma 1

