## Supplemental Proof 1 for "Effective Matrix Designs for COVID-19 Group Testing"

### PROOF OF LEMMA 1

DAVID BRUST AND JOHANNES J. BRUST

#### Proof of Lemma 1 ($d = 2$, and $q = p$ (prime))

Designs with $d = 2$ and $q = p$ (prime order), use modulus arithmetic and lines and may therefore be regarded to be simpler than more general polynomial constructions. Thus we provide a proof of Lemma 1 in this situation in order to motivate the underlying computations.

Similar like in the main text, suppose that $1 \leq k < p$ and let $j \neq i$. Select an arbitrary pair of samples $\{(x_i, y_{x_i}); (x_j, y_{x_j})\}$, computed using, say, $a$ and $b$. Consider the different slope $\overline{a} \neq a$ and intercept $\underline{b} \neq b$. There are four possibilities to generate pools in PP, and consequently pairs of samples other than a base pair

| | |
|---|---|
| Base: Slope and intercept given | $a, b$ with $\{(x_i, y_{x_i}); (x_j, y_{x_j})\}$ |
| Case 1: Slope differs | $\overline{a}, b$ with $\{(x_i, \overline{y}_{x_i}); (x_j, \overline{y}_{x_j})\}$ |
| Case 2: Intercept differs | $a, \underline{b}$ with $\{(x_i, \underline{y}_{x_i}); (x_j, \underline{y}_{x_j})\}$ |
| Case 3: Slope and intercept differ | $\overline{a}, \underline{b}$ with $\{(x_i, \underline{\overline{y}}_i); (x_j, \underline{\overline{y}}_j)\}$ |
| Case 4: Slope at infinity | $\overline{a} = q$ with $\{(x_i^\infty, y_i^\infty); (x_j^\infty, y_j^\infty)\}$ |

Next we analyze how the pairs in each of the four cases compare to the base pair $\{(x_i, y_{x_i}); (x_j, y_{x_j})\}$. We use $y^+ \in \{\overline{y}, \underline{y}, \underline{\overline{y}}\}$ to represent a $y$ value from one of the Cases 1–3. Note that the conditions for having different pairs of samples in these cases are

$$\text{if } y_{x_i} = y^+_{x_i} \text{ then } y_{x_j} \neq y^+_{x_j}, \quad \text{or} \quad \text{if } y_{x_j} = y^+_{x_j} \text{ then } y_{x_i} \neq y^+_{x_i} \tag{1}$$

For $d = 2$ the lines $y = (ax+b) \bmod p$ generate the different pools, which we use to evaluate condition (1). Moreover, with $d = 2$, the indices simplify to $i_{d-1} = i_1 = i$ and $x_i = i$, $x_j = j$.

*Proof.* Case 1: Only the slope $\overline{a}$ differs and we use the notation $y^+ = \overline{y}$. From the equation of a line and from (1)

$$y_{x_k} - \overline{y}_{x_k} = 0 \quad \text{implies} \quad ((a - \overline{a})x_k) \bmod p = 0 \quad \text{and} \quad x_k = 0, \quad k \in \{i, j\}$$

Since $x_k = k \neq x_t = t$ for $t \neq k \in \{i, j\}$ we see that

$$\text{if } y_{x_k} = \overline{y}_{x_k} \text{ then} \quad x_k = 0,\ x_t > 0, \quad y_{x_t} - \overline{y}_{x_t} = ((a - \overline{a})x_t) \bmod p \neq 0 \quad t \neq k \in \{i, j\}$$

Therefore, we conclude that $y_{x_j} \neq \overline{y}_{x_j}$ when $y_{x_i} = \overline{y}_{x_i}$ and similarly for the reverse $y_{x_i} \neq \overline{y}_{x_i}$ when $y_{x_j} = \overline{y}_{x_j}$. This means that pairs of samples do not co-occur when pools are generated in case 1.

Case 2: Only the intercept $\underline{b}$ differs and $y^+ = \underline{y}$. Note that

$$y_{x_k} - \underline{y}_{x_k} = (b - \underline{b}) \bmod p \neq 0, \quad k \in \{i, j\}$$

Therefore $y_{x_i} \neq \underline{y}_{x_i}$ and $y_{x_j} \neq \underline{y}_{x_j}$ and all samples and corresponding pairs are different in this case.

Case 3: Both slope $\overline{a}$ and intercept $\underline{b}$ differ and we denote $y^+ = \underline{\overline{y}}$. Using condition (1) one finds that

$$y_{x_k} - \underline{\overline{y}}_{x_k} = 0 \quad \text{implies} \quad ((a - \overline{a})x_k + (b - \underline{b}))\text{mod } p = 0, \quad k \in \{i, j\}$$

which means that

$$y_{x_k} - \underline{\overline{y}}_{x_k} = 0 \quad \text{implies} \quad ((a - \overline{a})x_k)\text{mod } p = -(b - \underline{b}))\text{mod } p, \quad k \in \{i, j\}$$

Therefore, for $k = x_k \neq x_t = t$ when $t \neq k \in \{i, j\}$ it holds that

$$\begin{aligned} \text{if } y_{x_k} = \underline{\overline{y}}_{x_k} \text{ then } y_{x_t} - \underline{\overline{y}}_{x_t} &= ((a - \overline{a})x_t + (b - \underline{b}))\text{mod } p \\ &= ((a - \overline{a})x_t - (a - \overline{a})x_k)\text{mod } p \\ &= ((a - \overline{a})(x_t - x_k))\text{mod } p \\ &\neq 0, \quad k \in \{i, j\} \end{aligned}$$

The final equality holds because $a \neq \overline{a}$ and $x_t \neq x_k$, $t \neq k \in \{i, j\}$ by definition.

Case 4: With slope of "infinity" $\in \mathbb{F}_p$ a pool of samples is computed with a special formula. Specifically, a pool (with $d = 2$) with this slope is defined to have $x_i^\infty = \sum_{l=2}^{2-1} p^{d-1-l} i_l + q^{2-2} b = b = x_j^\infty = b$ and

$$\{(x_i^\infty, y_i^\infty); (x_j^\infty, y_j^\infty)\} = \{(b, y_i^\infty); (b, y_j^\infty)\}.$$

Since $x_k \neq x_t$ for $k \neq t \in \{i, j\}$ (by definition) we note that if $x_k = b = x_k^\infty$ and $y_{x_k} = y_k^\infty$ then $x_t \neq x_t^\infty = x_k = b$. Thus no pairs occur more than once with the pools from slopes $a < q$ and $a = q$. Therefore, we conclude from the four cases that pairs occur jointly $d - 1 = 1$ times for all slopes $a \leq p$. $\square$
